# Adaptation of the Arizona Cognitive Task Battery for use with the Ts65Dn Mouse Model (*Mus musculus*) of Down syndrome

**DOI:** 10.1101/061754

**Authors:** Michael R. Hunsaker, Genevieve K. Smith, Raymond P. Kesner

**Author notes:** Current address: Special Education Department, Granite School District, 2500 S State Street, Salt Lake City, UT 84115. Please send correspondence and requests for offprint copies to: Raymond P. Kesner, Department of Psychology, University of Utah. The authors wish to acknowledge Dr. Julie R. Korenberg for providing access to facilities where these experiments were conducted. All authors declare they have no competing financial or professional interests.

## Abstract

We propose and validate a clear strategy to efficiently and comprehensively characterize neurobehavioral deficits in the Ts65Dn mouse model of Down syndrome. This novel approach uses neurocognitive theory to design and select behavioral tasks that test specific hypotheses concerning the results of Down syndrome. In this manuscript we model in Ts65Dn mice the Arizona Cognitive Task Battery used to study human populations with Down syndrome. We observed specific deficits for spatial memory, impaired long-term memory for visual objects, acquisition and reversal of motor responses, reduced motor dexterity, and impaired adaptive function as measured by nesting and anxiety tasks. The Ts65Dn mice showed intact temporal ordering, novelty detection, and visual object recognition with short delays. These results phenocopy the performance of participants with Down syndrome on the Arizona Cognitive Task Battery. This approach extends the utility of mouse models of Down syndrome by integrating the expertise of clinical neurology and cognitive neuroscience into the mouse behavioral laboratory. Further, by directly emphasizing the reciprocal translation of research between human disease states and the associated mouse models, we demonstrate that it is possible for both groups to mutually inform each others’ research to more efficiently generate hypotheses and elucidate treatment strategies.

## Introduction

One reason we propose underlying the lack of direct applicability of mouse model research for improving the quality of life of people with developmental disabilities is an unfortunate focus on gross phenotypes that may be either at best secondary to the mutation or result from mouse-unique factors that do not scale evolutionarily to humans. Stated more colloquially, it is much easier to cure disease in mice than to translate the murine research into actually curing human disease. The same general paradigm is prevalent in research into sequelae resultant to neurodevelopmental/neurodegenerative genetic diseases. One solution to this difficulty is to specifically design behavioral paradigms to test in mice what is being tested in human research participants. This process is called behavioral or neurocognitive endophenotyping (Gottesman & Gould, 2003; Hunsaker, 2012a, 2012b; Simon, 2008).

There is a clear difference between identifying a behavioral phenotype and identifying a behavioral endophenotype. This difference is that to evaluate a behavioral phenotype, the researcher need only look for a difference in behavior among a homogeneous group of mutant mice relative to littermate or strain-matched control group. This main effect is then used as evidence for some kind of behavioral impairment. This process is akin to using the same battery of standardized neuropsychological tests to evaluate the behavioral consequences of number of different genetic disorders and then trying to make inferences about what are the specific profiles of strengths and weaknesses unique to each disorder. In contrast, to evaluate a behavioral endophenotype in the same mice, there is a requirement that any behavioral phenotype predictably scale across some measure: Usually such factors include age, genetic dosage in situations of polymorphic mutations or chromosomal aneuploidy, or some other experimentally controlled factor that is altered parametrically (*e.g.*, stress, environmental toxicant exposure, etc.). This process is similar to how experimental psychology or cognitive neuroscience approaches to studying the behavior of populations carrying genetic mutations. That is, an approach that emphasizes using hypothesis driven tests that have been designed to evaluate hypothesized effects within the population being studied, irrespective to performance of other populations.

The importance of finding a behavioral endophenotype is that if there is a predictable relationship among cognitive performance and gene expression, it can be assumed that the genetic mutation alters behavioral output; and subsequently, some sort of relationship between the two exists. Such a finding not only provides a wealth of information that helps the researcher design future experiments, but also data that are useful as outcome measures for studies of intervention that alter or even potentially mitigate some negative impact of the mutation. If there is a more complex relationship wherein age appears to modulate the relationship between the mutation and behavioral output, then those data serve not only as outcome measures, but if well enough understood, could be potentially useful to define risk prodromes to predict future symptomatology or disease progression (*[cf.],*Gottesman and Gould (2003)).

As a scientific community, we have been able to identify and provide cures for a wide range mouse models of genetic disorders (*i.e.*, Down Syndrome), but to date these cures have not proven particularly useful for ameliorating symptoms of human genetic disease: often failing or providing only marginal effects during early phase clinical trials. Elucidating behavioral or neurocognitive endophenotypes using tasks designed to test specific disease-related hypotheses is one proposed solution to mitigate this lack of efficacy in the mouse model.

For these, as well as many other reasons, research into schizophrenia has forced the field to changed their general approach, and emphasized an endophenotyping approach in the study of prodromal states associated with schizophrenia onset and symptom progression (*e.g.*, focusing research on longitudinal analyses of 22q11.2 deletion populations rather than on *de novo* schizophrenia cases of unknown or poorly understood genetic origin; Gottesman and Gould (2003), Karayiorgou, Simon, and Gogos (2010), Simon (2008)). By focusing on factors that scale with disease or symptom severity, researchers have been able to understand far more about schizophrenia and what may underlie symptom progression than they would otherwise have been able using a standardized, neuropsychological phenotyping approach.

Mouse models often demonstrate phenotypes that are not specifically associated with any genetic disorder in particular, but are more aptly described as shared clinical phenotypes that similarly present across a wide array of disorders (*e.g.*, global learning and memory deficits, dementia, anxiety, depression). The interpretation of such inconclusive findings is often that the mouse model fails to recapitulate the phenotypes observed in patients. Unfortunately, these types of findings are analogous to inconsistent findings in clinical populations when standardized neuropsychological tests are administered – many different populations show very similar deficits despite nonoverlapping genetic or developmental disorders. Such inconsistencies often renders behavioral research into developmental or psychiatric disorders frustrating and such anomalous findings mask the differences that do exist. Hunsaker (2012a, 2012b, 2013), Simon (2008) proposed that inconsistent behavioral results observed in clinical populations as well as mouse models do not infer the lack of cognitive impairments, but rather these "null" data reflect the often startling insensitivity of the behavioral tasks commonly employed.

In situations where, based on standardized behavioral tasks, mouse models do not appear to specifically model clinical phenotypes observed in patient populations, one strategy is to evaluate intermediate- or endophenotypes associated specifically with the genetic mutation and subserved by neuroanatomical structures disrupted by the mutation. A similar process applies to studies of human clinical populations when standardized tests fail to uncover phenotypes that are present, but only manifest at a subclinical level. Endophenotypes are collections of quantitative traits hypothesized to represent risk for genetic disorders at more biologically (and empirically) tractable levels than the full clinical phenotype; which often contains little more than profound deficits shared across various genetic disorders.

A behavioral endophenotyping approach facilitates the identification of behavioral deficits that are clearly associated with both the specific genetic mutation and the pathological features observed in the clinical populations being modeled – and more importantly with the pathological/clinical features unique to the population being modeled. When designed to evaluate such disease-specific hypotheses, behavioral endophenotypes model quantitative patterns of behavioral deficits that scale with the size and/or severity of the genetic mutation.

The behavioral endophenotyping process deviates from the currently accepted method for determining behavioral phenotypes. The currently accepted method to determine phenotypes in clinical populations and mouse models is to use behavioral tasks that were designed without prior consideration of the pathology and clinical features present in the population. Far too often an approach such as this is not sufficiently sensitive to characterize the gene-brain-behavior interactions that underlie disease pathogenesis. In contrast with the currently utilized approach, behavioral endophenotyping emphasizes the use of behavioral paradigms that were developed to specifically evaluate a priori hypotheses concerning the alterations to nominal gene-brain-behavior interactions identified (or proposed to exist) in a given patient population using carefully selected tasks designed to identify unique phenotypes within each model; and thus are more capable of characterizing the neurocognitive consequences of the specific gene mutations underlying the genetic disorder.

In order to design a battery of behavioral/neurocognitive tasks that could be presented to individuals with Down syndrome across a wide age range in a single testing session, Edgin et al. (2010) developed and validated the Arizona Cognitive Task Battery (ACTB). What makes this battery different than others that are available at present (*e.g.*, Cambridge Neuropsychological Testing Automated Battery (CANTAB)) is that the ACTB has been developed to keep the following issues in mind: 1) when one studies a population with a neurodevelopmental disease, particularly a chromosomal aneuploidy, there is a very real possibility of floor effects confounding analyses of behavioral or cognitive task performance. 2) Additionally, individuals with Down syndrome show language deficits, limiting the tasks that can be used to test cognitive function without a language confound. 3) Finally, and perhaps most importantly, the ACTB was developed with the goal of maximizing the sensitivity to identify effects that are present in Down syndrome.

The IQ in Down syndrome is typically moderately to severely intellectually disabled range (*i.e.*, IQ = 25-55) and mental age rarely moves beyond 8 years. Paradoxically, it has been suggested that early on, Down syndrome only presents with a mild to moderate intellectual disability (*i.e.*, 55-70), but with age the IQ drops as mental age no longer increases with chronological age (Edgin et al., 2010; Virji-Babul, Kerns, Zhou, Kapur, & Shiffrar, 2006).

It has been hypothesized that visual-spatial abilities appear to be normal in Down syndrome. However, this appears to be something of an artifact when visual-spatial memory is directly compared to auditory and verbal performance. In tests specifically assessing visual and spatial abilities in Down syndrome, there is a clear deficit relative to typically developing or age matched control populations (Edgin et al., 2010; Edgin, Mason, Spano, Fernandez, & Nadel, 2012; Pennington, Moon, Edgin, Stedron, & Nadel, 2003).

Within the memory domain, Down syndrome results in deficits for digit or word span as well as general memory deficits with long delays prior to recall. Working memory, specifically verbal working memory, is disrupted in Down syndrome (Edgin, Spano, Kawa, & Nadel, 2014; Pennington et al., 2003; Stedron, Sahni, & Munakata, 2005; Vicari, Bellucci, & Carlesimo, 2005). For visual and spatial memory, it appears that Down syndrome results in specific memory deficits when memory span is increased (Carretti & Lanfranchi, 2010; Lanfranchi, Carretti, Spano, & Cornoldi, 2009; Silvia Lanfranchi, Cornoldi, Vianello, & Conners, 2004). Again, as suggested by the language deficits, it has been shown that individuals with Down syndrome have greater impairments for verbal than visual-spatial span. Down syndrome also results in long-term memory deficits (Pennington et al., 2003; Vicari, 2006).

Despite these memory deficits, implicit memory and perceptual priming appear to be normal (Pennington et al., 2003; Vicari, 2006). This pattern suggests that there is an explicit memory deficit in Down syndrome, meaning that when memory requires temporal or spatial processing, there is a deficit. This has implicated hippocampus and medial temporal lobe function in Down syndrome pathology, as well as the prefrontal cortex for working memory. Implicit memory, dependent upon different brain areas (*e.g.*, parietal cortex), appears to be spared, if not slightly facilitated in Down syndrome compared to other cognitive domains (*i.e.*, word stem or perceptual priming tasks).

It has been shown that motor development in Down syndrome is slower than age and mental age matched peers. Intriguingly, early motor markers like rolling and sitting up have been shown to be only very subtly slowed in Down syndrome, but crawling and walking has been shown to be more dramatically delayed. Despite this delay, it does appear that children with Down syndrome develop through the same milestones as typically developing children, these milestones just occur dramatically later in development. Motor skill development appear to show the same developmental delays as these early markers of motor abilities (Connolly & Michael, 1986; Frith & Frith, 1974; Gemus et al., 2002; Rast & Harris, 1985; Vicari, 2006; Virji-Babul et al., 2006).

To date, the majority of behavioral assays used to test the behavioral phenotype of the mouse models of Down syndrome have focused on spatial memory. More specifically, focus has been placed on the Morris water maze test of spatial memory (Escorihuela et al., 1995; Reeves et al., 1995; Sago et al., 1998). Later experiments have focused on novel object recognition at short and long delays as a proxy for general memory deficits observed across wide range of mouse disease models (Faizi et al., 2011). As a measure of executive function or rostral cortical function, spontaneous alternation has been used (A. M. Kleschevnikov et al., 2012; A. M. Kleschevnikov et al., 2004). The majority of motor tests use the rotarod or locomotor behavior in an open field as the primary measure (Faizi et al., 2011).

In this study we propose and then evaluate a clear strategy to efficiently and comprehensively characterize neurobehavioral deficits in the Ts65Dn mouse model of Down syndrome by developing a mouse variant of the Arizona Cognitive Task Battery (Mouse Cognitive Task Battery; mCTB). This approach uses neurocognitive theory to design and select behavioral tasks that test specific hypotheses concerning the genetic disorder being studied-specifically those proposed as part of the Arizona Cognitive Task Battery (ACTB) used to study human populations with Down syndrome (Edgin et al., 2010; Hunsaker, 2012a).

This approach specifically relies on known anatomical data regarding human and mouse model brain function as important considerations in task design and selection, similar to the ACTB (Edgin et al., 2010). This approach extends the utility of mouse models by integrating the expertise of clinical neurology and cognitive neuroscience into the mouse behavioral laboratory. Further, by directly emphasizing the reciprocal translation of research between human disease states and the associated mouse models, we demonstrate that it is possible for both groups to mutually inform each others’ research to more efficiently generate hypotheses and elucidate treatment strategies (*cf.*, Hunsaker, 2012a, 2016).

## Materials and Methods

### Animals

In this study, 10 segmentally trisomic Ts(1716)65Dn (Ts65Dn) male mice and 10 age-matched wildtype littermates were obtained from Jackson Laboratories (Bar Harbor, ME) and tested at 5-7 months of age, weighing 33 +/- 5.2g (SD). Ten mice per group was chosen as the minimum number of mice required to obtain a reliable behavioral result based on a predictive power analysis using data from similar tasks reported by previous studies using the CGG Knock-In and *Fmr1* knockout mouse models *cf.*, Hunsaker (2012a, 2013). The Ts65Dn/DnJ stock, commercially available from Jackson Laboratory (B6EiC3Sn.BLiA-Ts(1716)65Dn/DnJ), is homozygous for the wildtype allele for retinal degeneration. The stock is maintained by repeated backcrossing of Ts65Dn females to B6EiC3H F1 hybrid males derived from a new congenic strain of C3H mice. This new congenic strain (C3Sn.BLiA-Pde6b+) lacks the blindness causing recessive mutant allele. Animals were kept on a 12-h light/dark cycle, in a temperature and humidity controlled environment with *ad libitum* access to food and water. During no point in experimentation was food deprivation used. Care was taken to assure mice showed motivation to seek sucrose pellet rewards. All behavioral tests were conducted during the light portion of the cycle (06:00-18:00). Mice were housed in same-genotype groups of 2-3 per cage. Animal care and experimental testing procedures conformed to NIH, IACUC, and AALAC standards and protocols.

### Experimental Design for Behavioral Testing

The week prior to testing, all animals were handled daily for 15 min sessions and given an opportunity to habituate to a clear and red apparatus for at least 15 min each and acclimate to sucrose pellet rewards. It was verified that prior to the end of this training period that mice consumed sucrose pellets as soon as placed on the apparatus. Behavioral tasks emphasizing exploratory behaviors were presented in a pseudo-randomized order between mice (randomized within the Ts65Dn mice and a 2N wildtype littermate was yoked to a given Ts65Dn mouse to account for any potential task order effects), followed by spontaneous alternation and motor tasks, then response and reversal learning tasks. The 2N wildtype mice were the same age (within 15 days of age) as the Ts65Dn mice.

After these tasks, mice received training on the cheeseboard, and then finally were presented with test designed to evaluate quality of life/adaptive functional measures to reduce the influence of any anxiety measures on later task performance.

To specifically isolate the contribution of spatial and non spatial cues to task performance, behavioral tasks were run two times, once in a clear box and many extra maze cues, and a second time in a red box without extra maze cues (Dees & Kesner, 2013). This was done because Smith, Kesner, and Korenberg (2014) noticed that there was a pattern of deficits in Ts65Dn mice that were better explained by the mice having access to the extra-maze context than by any specific memory process. As such, they ran every experiment twice, one time using a clear box that allowed access to extra-maze cues and another time in a red box that blocked the view of the extra maze cues. They found that visual object recognition deficits at a 1 hour delay were seen in the clear box experiment, whereas experiments in the red box showed intact visual object memory at a 1 hour delay. They attributed this effect to extra-maze or distal context interfering with the visual object recognition due to interference. Experiments in rats exploring the same effect revealed similar results, and further unpacked the neural correlates of this effect Dees and Kesner (2013). The rationale for this procedure comes from work reported by Smith et al. (2014) in Ts65Dn mice and Edgin et al. (2014) in children with Down syndrome showing that context is particularly influential during object recognition tasks in children with Down syndrome relative to typically developing children. In other words, children with Down syndrome are particularly susceptible to memory interference during cognitive tasks.

For every experiment a novel set of objects were used, such that no mouse ever encountered the same object during different experiments. At the end of every experiment, 95% ethanol was used to reduce and spread olfactory cues and prevent odor effects impacting future task performance.

### Tests of Spatial Attribute

#### Spatial Navigation using Cheeseboard

Apparatus: A white, circular Plexiglas platform with a series of 2 cm diameter holes centered every 5 cm was used as the cheeseboard apparatus. The apparatus was placed approximately 1.5 m off the ground in a space surrounded by extra maze, distal cues to provide a rich spatial context to guide mouse navigation. Paths taken by the mice were recorded by an overhead camera and analyzed using Noldus EthoVision software.

Method: Each mouse was habituated to the cheeseboard for 30 min the day prior to experimentation with banana flavored sucrose pellets distributed in each hole (Bio-Serv, #F07257). All mice consumed sucrose pellets and showed a random foraging pattern prior to beginning of training. At the beginning of each trial, a single sucrose reward pellet was placed in one of the holes of the cheeseboard (located within the midpoint of the North-East, North-West, South-East or South-West quadrant). A mouse was then released at one of the cardinal points (*e.g.*, North, South, East, or West at the edge of the cheeseboard) as latency in seconds and distance in centimeters traveled to locate and consume the reward was recorded. Each day, the mouse received a trial from each of the four cardinal directions (order randomized between mice and between days within mice). There were 5 minutes separating each trial for each mouse. After the fourth day of training, the mice were given a probe trial wherein there was no reward. The search patterns of the mice were evaluated. This protocol was modified from the original rat protocol (Kesner, Farnsworth, & DiMattia, 1989) for mice after experiments reported by Lopez, Hauser, Feldon, Gargiulo, and Yee (2010).

#### Metric/Coordinate Processing

Apparatus: The apparatus for these experiments consisted of a large Plexiglas box 40 cm wide by 40 cm deep with clear walls 40 cm in height and a dark gray floor. An inset made of translucent red Plexiglas 39 cm in width x 39 cm in height was constructed for easy insertion and removal from the original clear box, therefore enabling the experimenter to block distal cues in the testing environment when desired. The box was placed on a circular white table 1 m in diameter. Four distinct two-dimensional black and white cues were placed 30 cm away from each side of the box (methods after Smith et al. (2014)). Exploration was recorded with an overhead video camera and the duration of exploration was measured with a stopwatch. Proximal objects were made from various washable, non-porous materials (plastic, metal, glass, etc.), ranging 2-7 cm in height and had various color, pattern, and textures to ensure each object was visually distinct. New objects were used between experiments so mice were never exposed to the same object during different experiments. To prevent use of olfactory cues to guide behavior, the boxes and objects were disinfected and deodorized with a sterilizing cleaning agent after each use. The mouse was presented with entirely novel object sets for every experiment. All locomotor activity was collected by the Noldus EthoVision software calibrated to measure to the nearest cm (Noldus USA, North Carolina).

**Table 1.** Comparison of Arizona Cognitive Task Battery (ACTB) and Mouse Variant Reported in this Manuscript (mCTB). The mCTB was designed to model as many of the functions as the ACTB was designed to tests in humans. Cognitive deficits summarized in the table phenocopy the effects seen in Down syndrome on the ACTB or subsequent follow-up studies (Edgin et al., 2010; Edgin, Mason, Spano, Fernandez, & Nadel, 2012). Similarly, the performance of Ts65Dn mice on the mCTB recapitulates intact cognitive function seen in participants with Down syndrome when tested using the ACTB

| Domain/Test in ACTB | Abilities Assessed | Analogous Task in mCTB | Ts65Dn performance |
| --- | --- | --- | --- |
| ***Benchmark, General Cognitive Ability*** |  |  |  |
| KBIT-II Verbal Subscale | Receptive and Productive Language | *not modeled* | n/a |
| KBIT-II Nonverbal Subscale | Problem Solving | *not modeled* | n/a |
| Scales of Independent Behavior-Revised (SIB-R) | Adaptive Function | Nesting, Neophobia | **deficits for Adaptive Function** |
| CANTAB Spatial Span | Immediate Memory for Spatiotemporal Information | Temporal Order for Visual Objects | **no deficits for Temporal Order** |
| ***Prefrontal Cortex - Executive Function, Response Attribute*** |  |  |  |
| Modified Dots Task | Inhibitory Control and Working Memory | Spontaneous Alternation | **deficits for Spontaneous Alternation** |
| CANTAB IED | Set Shifting | Rule Response Learning, Rule Reversal Learning | **deficits for Response Learning, deficits for Reversal Learning** |
| ***Medial Temporal Lobe - Spatial Attribute*** |  |  |  |
| CANTAB PALS | Spatial Associative Memory | Location Recognition | **deficits for Location Recognition** |
| Virtual Water Maze | Spatial Memory/Navigation | Dry Land Water Maze (Cheeseboard) | **deficits for Acquisition and Retrieval of Spatial Navigation** |
| *not evaluated* | Spatial Relationships | Coordinate, Categorical | **deficits for Coordinate task, no deficits for Categorical task** |
| ***Medial Temporal Lobe - Temporal Attribute*** |  |  |  |
| *not evaluated* | Temporal Processing/Sequence Learning | Temporal Order for Visual Objects | **no deficits for Temporal Order** |
| ***Medial Temporal Lobe - Sensory/Perceptual Attribute*** |  |  |  |
| *not evaluated* | Object Recognition | Feature Ambiguity, Object Recognition, Novel Object Detection | **No deficits at 1 hour delay, deficits at 24 hour delay** |
| ***Cerebellum - Motor Function*** |  |  |  |
| Finger Sequencing Task | Motor Sequencing | Capellini Handling | **deficits for Motor Sequencing** |
| NEPSY Visuomotor Precision | Visuomotor Tracking/Hand-Eye Coordination | Parallel Rung Walk, Capellini Handling | **deficits for Motor Coordination** |
| CANTAB SRT | Motor Response Time/Attention | *not modeled* | n/a |

Method: Each mouse had previously been habituated to clear and red experimental boxes. For the metric/coordinate processing test (Hunsaker, 2012a, 2013; Hunsaker, Kim, Willemsen, & Berman, 2012; Hunsaker, Wenzel, Willemsen, & Berman, 2009; Kesner et al., 2014; Smith et al., 2014), two objects were placed in the box separated by 25 cm (from inner edges) and mice were allowed to explore the objects for 15 minutes. After a 5 min interval during which the mice were covered by an opaque, heavy cup, the objects were moved closer together to an 8 cm separation and the mouse was allowed to explore for 5 min. This procedure was carried out in the clear box that allowed the mouse to see the extra-maze, distal cues as well as in the red box that blocked the ability of the mouse to see these cues (Dees & Kesner, 2013; Smith et al., 2014). Exploration during the last 5 min of habituation and during the 5 min test session were converted into a ratio value ranging [-1,1] to control for overall exploration. As such, a ratio value approaching -1 is interpreted as the mouse showing continued habituation and thus not noticing the change. A ratio value approaching 1 suggest the mouse dramatically explored the change.

#### Topological/Categorical Processing

Apparatus: This experiment used the same apparatus as the Metric/Coordinate experiment. A similar ratio value was computed as a dependent measure.

Method: Each mouse had previously been habituated to clear and red experimental boxes. For the topological/categorical processing test (Hunsaker, 2012a, 2013; Hunsaker et al., 2012; Hunsaker et al., 2009; Kesner et al., 2014; Lee et al., 2009; Smith et al., 2014), four objects were placed in a square in the box separated by 25 cm (from inner edges) and mice were allowed to explore the objects for 15 minutes. After a 5 min interval during which the mice were covered by a heavy cup, the front two objects were transposed, and the mouse was allowed to explore for 5 min. This procedure was carried out in the clear box that allowed the mouse to see the extra-maze, distal cues as well as in the red box that blocked the ability of the mouse to see these cues. Exploration during the last 5 min of habituation and during the 5 min test session were converted into a ratio value ranging [-1,1] to control for overall exploration. As such, a ratio value approaching -1 is interpreted as the mouse showing continued habituation and thus not noticing the change. A ratio value approaching 1 suggest the mouse dramatically explored the change in the object’s spatial location and/or distance from each other.

#### Spatial Location Recognition

Apparatus: This experiment used the same apparatus as the Metric/Coordinate experiment. A similar ratio value was computed as a dependent measure using exploration data.

Method: Each mouse had previously been habituated to clear and red experimental boxes. For the location recognition test (Smith et al., 2014), two objects were placed in the box separated by 25 cm (from inner edges) and mice were allowed to explore the objects for 15 minutes. After a 5 min interval during which the mice were covered by a heavy cup, one of the objects was moved at a diagonal to a new location (still 25 cm separation between the two objects), and the mouse was allowed to explore for 5 min. This procedure was carried out in the clear box that allowed the mouse to see the extra-maze, distal cues as well as in the red box that blocked the ability of the mouse to see these cues. Exploration during the last 5 min of habituation and during the 5 min test session were converted into a ratio value ranging [-1,1] to control for overall exploration. As such, a ratio value approaching -1 is interpreted as the mouse showing continued habituation and thus not noticing the change. A ratio value approaching 1 suggest the mouse dramatically explored the change in which object occupied which spatial location.

### Tests of Temporal Attribute

#### Temporal Ordering for Visual Objects

Apparatus: This experiment used the same apparatus as the Metric/Coordinate experiment. A similar ratio value was computed as a dependent measure.

Method: During session 1, two identical copies of a first object (object 1) were placed at the ends of the box 2.5 cm from the end walls and centered between the long walls (Hunsaker, 2013; Hunsaker, Goodrich-Hunsaker, Willemsen, & Berman, 2010; Hunsaker et al., 2012). The mouse was placed in the center of the box facing away from both objects. The mouse was given 5 min to freely explore the objects. After 5 min, the mouse was removed to a small holding cup for 5 min. During this time, the first objects were replaced with two duplicates of a second object (Object 2). For Session 2, the mouse was again placed in the apparatus and allowed to explore. After 5 min, the mouse was removed to the holding cup for 5 min and the objects were replaced with two duplicates of a third object (Object 3). For Session 3, the mouse was given 5 min to explore. After 5 min, the mouse was removed into a small cup for 5 min and an unused copy of the first and an unused copy of the third object were placed into the box. The mouse was again placed into the box and allowed to explore the two objects (*i.e.*, Objects 1 and 3) during a 5 min test session. This procedure was carried out in the clear box that allowed the mouse to see the extra-maze, distal cues as well as in the red box that blocked the ability of the mouse to see these cues. Exploration of each object during the test session were converted into a ratio value ranging [-1,1] to control for overall exploration. As such, a ratio value approaching -1 is interpreted as the mouse showing an absolute preference for the third over the first object. A ratio value approaching 1 suggest the mouse strongly explored the first over the third object.

#### Temporal Order Control - Novelty Detection for Visual Objects

Apparatus: This experiment used the same apparatus as the Metric/Coordinate experiment. A similar ratio value was computed as a dependent measure.

Method: In addition to reflecting impaired temporal ordering, increased exploration of the first object over the third could also be interpreted as being due to difficulty in remembering the first object prior to the test session (Hunsaker, 2012a, 2013; Hunsaker et al., 2010). To minimize and control for such general memory deficits, a novelty detection of visual objects task was performed. Briefly, on a different day mice received three sessions during which they were allowed to explore three novel sets of objects (Objects 4, 5, 6) similarly to the temporal ordering tasks. During the test session, the first object and a novel fourth object (Object 7) were presented and the mice were allowed 5 min to explore. This procedure was carried out in the clear box that allowed the mouse to see the extra-maze, distal cues as well as in the red box that blocked the ability of the mouse to see these cues (*cf.*, Dees and Kesner, 2013; Smith et al., 2014). Exploration of each object during the test session were converted into a ratio value ranging [-1,1] to control for overall exploration. As such, a ratio value approaching -1 is interpreted as the mouse showing an absolute preference for the familiar over the novel object. A ratio value approaching 1 suggest the mouse strongly explored the novel over the familiar object.

### Sensory/Perceptual Attribute

#### Feature Ambiguity

Apparatus: This experiment used the same apparatus as the Metric/Coordinate experiment. A similar ratio value was computed as a dependent measure.

Method: Each mouse had previously been habituated to clear and red experimental boxes. For the configural recognition condition (Bartko, Winters, Cowell, Saksida, & Bussey, 2007; Bussey, Saksida, & Murray, 2002, 2006; Smith et al., 2014), mice were placed for 15 min in the red box containing two compound objects, A-B and C-D, separated by 15 cm. Following a 5 min delay under a heavy cup, the mouse underwent a 5-min Test Phase in which one object from the Study Phase remained the same (A-B) and the other compound object is created from one component of each of the previous familiar objects, (*e.g.*, A-D). That is, the "novel" object (A-D) was composed of the same elements, but rearranged into a novel configuration. Therefore, the object is "novel" by virtue of its configuration, not by its elements, each of which was present in one of the original compound stimuli. Exploration of each compound object was scored as a single unit. Exploration during the last 5 min of habituation and during the 5 min test session were converted into a ratio value ranging [-1,1] to control for overall exploration. As such, a ratio value approaching -1 is interpreted as the mouse showing continued habituation and thus not noticing the change. A ratio value approaching 1 suggest the mouse dramatically explored the change.

#### Feature Ambiguity Control - Novelty Detection for Configuration of Objects

Apparatus: This experiment used the same apparatus as the Metric/Coordinate experiment. A similar ratio value was computed as a dependent measure.

Method: Each mouse had previously been habituated to clear and red experimental boxes. For the configural recognition condition (Bartko et al., 2007; Bussey et al., 2002, 2006; Smith et al., 2014), mice were placed for 15 min in the red box containing two compound objects, A-B and C-D, separated by 15 cm. Following a 5 min delay under a heavy cup, the mouse underwent a 5-min control task during which C-D was replaced by two never before seen objects (E-F) was also performed. This procedure was carried out in the clear box that allowed the mouse to see the extra-maze, distal cues as well as in the red box that blocked the ability of the mouse to see these cues. Exploration during the last 5 min of habituation and during the 5 min test session were converted into a ratio value ranging [-1,1] to control for overall exploration. As such, a ratio value approaching -1 is interpreted as the mouse showing continued habituation and thus not noticing the change. A ratio value approaching 1 suggest the mouse dramatically explored the change.

#### Object Recognition at 1 and 24 Hour Delays

Apparatus: This experiment used the same apparatus as the Metric/Coordinate experiment. A similar ratio value was computed as a dependent measure.

Method: Each mouse had previously been habituated to clear and red experimental boxes. For the object recognition test (Moore, Deshpande, Stinnett, Seasholtz, & Murphy, 2013; Smith et al., 2014), two objects were placed in the box separated by 25 cm (from inner edges) and mice were allowed to explore the objects for 15 minutes. After a 5 min interval during which the mice were covered by a heavy cup, one of the objects was replaced by a novel object that had never before been experienced by the mouse, and the mouse was allowed to explore for 5 min. This procedure was carried out in the clear box that allowed the mouse to see the extra-maze, distal cues as well as in the red box that blocked the ability of the mouse to see these cues. This procedure was carried out in each box separately for delays of 1 hour and 24 hours. Exploration during the last 5 min of habituation and during the 5 min test session were converted into a ratio value ranging [-1,1] to control for overall exploration. As such, a ratio value approaching -1 is interpreted as the mouse showing continued habituation and thus not noticing the change. a ratio value approaching 1 suggest the mouse dramatically explored the change.

### Tests of Executive Function

#### Spontaneous Alternation

Apparatus: For this experiment, a Y maze with each arm measuring 45 cm in length by 30 cm in height with a runway width of 6 cm was used. It was made from opaque gray Plexiglas to prevent the use of any extra-maze cues to guide behavioral performance. As this was a spontaneous alternation task, no rewards were provided at the end of the arms of the Y maze.

Method: Mice were placed in the stem of a Y maze and allowed to explore (Faizi et al., 2011; A. M. Kleschevnikov et al., 2012; A. M. Kleschevnikov et al., 2004). Whenever the mouse entered one of the arms of the Y maze with all four limbs their response was recorded. Upon reaching the end of the arm, the mouse was gently picked up and replaced in the stem of the Y maze. The number of times the mouse alternated (*i.e.*, did not repeat the previous turn), was recorded as an alternation.

#### Response Learning

Apparatus: For this experiment, a plus maze with each arm measuring 50 cm in length by 25 cm in height with a runway width of 8 cm was used. There was a 2 cm diameter depression at the end of the arms wherein a sucrose pellet was placed to reward a correct response. It was made from opaque gray Plexiglas to prevent the use of any extra-maze cues to guide behavioral performance. At any time the mouse was required to make a 90 degree turn to the right or left to make a choice. The remaining arm was blocked off using a gray Plexiglas block that fit snugly into the arms of the plus maze.

Method: Mice were placed in the stem of a plus maze with one of the arms blocked off (forming a T maze). Mice were given five trials to determine if there was any preference for one direction over the other. As no such preference was observed, mice were randomly assigned the rule to turn right or turn left. Mice received 20 trials per day for 4 days (Bissonette et al., 2008; Ragozzino, Detrick, & Kesner, 1999; Ragozzino, Ragozzino, Mizumori, & Kesner, 2002). Entry into an arm with all four limbs was recorded as a choice and mice were not allowed to self correct when they made mistakes. Upon reaching the end of the arm, the mouse was gently picked up and replaced in the stem of the plus maze.

#### Reversal Learning

Apparatus: This experiment is a continuation of the Response acquisition experiment and used the same apparatus. For this experiment, the previously rewarded arm was now unrewarded and the previously unrewarded arm was now rewarded by a sucrose pellet.

Method: The day after mice finished training on response learning, they received 80 trials of reversal training (Bissonette et al., 2008; Ragozzino et al., 1999; Ragozzino et al., 2002). This means that the turn the mice had just learned to make for reward was now incorrect, rather the mice had to make the opposite turn to receive reward. Upon reaching the end of the arm, the mouse was gently picked up and replaced in the stem of the plus maze. Number of previously correct choices made were recorded as errors and error type was evaluated as perseverative or regressive based on the work of Aggleton and Ragozzino (Ragozzino et al., 2002; E Clea Warburton, Baird, Morgan, Muir, & Aggleton, 2001; E. Warburton, Baird, Morgan, Muir, & Aggleton, 2000). Briefly, errors during trials 1-20 were considered perseverative errors (perseverating or inflexibly following a previously learned rule) and errors during trials 21-40 were considered regressive errors (regressing or returning to a previously learned rule). Additionally, a behavioral change point algorithm was used to define the point at which each mouse consistently switched their responses from the previously learned rule to the new rule. This was done after the work reported by Diep et al. (2012) by taking the derivative of the learning curve at each point and evaluating when the derivative significantly changed slope (analysis code available at http://www.github.com/mrhunsaker/Change_Point).

### Motor Function

#### Capellini Handling

Apparatus: For this experiment, a 250 mL Nalgene beaker was used as a testing environment to assist in video recording mouse behavior. A small mirror was set up behind the beaker and the camera was placed to capture a front and rear view of the mouse to record trials.

Method: Mice were habituated over a weekend with approximately 20-30 dried capellini pasta presented in their cages (Tennant et al., 2010). Each mouse was placed in a 250 mL beaker and given a 5 cm piece of dried capellini. Their behaviors while eating were recorded for an offline analysis of their motor behaviors. Their latency to finish each piece of pasta was recorded, as were abnormal behaviors including the mouse having its paws together while eating, losing contact with the pasta with one or both paws, and using the mouth to pull the pasta rather than using the digits to feed the pasta into the mouth.

#### Parallel Rung Walking

Apparatus: Mice were placed in a box measuring 15 cm wide by 15 cm deep by 45 cm tall with 1.5 mm diameter parallel rungs making up the floor. The rungs were designed with same spacing used by Hunsaker et al. (2011). However, as this was a box rather than a runway, locomotor activity was collected using the Noldus EthoVision software to evaluate any effects of locomotor activity on motor coordination.

Method:The mice were allowed to freely explore the box for 5 minutes (Cummings, Engesser-Cesar, Cadena, & Anderson, 2007; Farr, Liu, Colwell, Whishaw, & Metz, 2006; Hunsaker et al., 2011). The number of times a paw slipped through the parallel rod floor beyond the wrist or ankle, a "foot slip" error was recorded (protocol simplified after Farr et al. (2006)). Total number of steps was also recorded to be used as an adjustment factor in later analyses.

### Adaptive Function

#### Nesting Behaviors

Apparatus: A 10 cm long piece of 5 cm diameter PVC pipe capped at one end was used as the apparatus. Sawdust similar to that used as mouse bedding was used as a nesting substrate.

Method: Sawdust was used to fill a 10 cm long piece of 5 cm diameter PVC pipe that was capped at one end (dry fit, no glue was used). This pipe was placed in a cage with each mouse and the latency to contact the sawdust in the pipe, the latency to start digging in the sawdust, and the latency to finalize the nest were recorded (Filali & Lalonde, 2009).

#### Neophobia

Apparatus: The home cage of the mouse, a 35 cm diameter metal platter, and a novel white Plexiglas box measuring 15 cm in all dimension were used to assess neophagia.

Method: Mice were given three neophobia tests (specifically hyperneophagia tests) based on the work of Bannerman et al. (2002). The first test was in each mouse’s home cage. Each mouse was provided a food they had never encountered (Cheerios cereal) and the latency for the mouse to take the first bite was recorded. The second test was each mouse was placed on a large platter in a bright area in the testing room and the latency for the mouse to take a bite from a reward pellet (familiar food) was recorded. The final test consisted of each mouse being placed in a novel white box and fed a Cheerio that had been stored in a sealed container filled with thyme overnight, resulting in a novel food (Vale-Martinez, Baxter, & Eichenbaum, 2002). Again, latency for the mouse to take the first bite was recorded.

### Statistical Methods

#### Dependent Measures and Data Visualization

For the Dry Land Water Maze on the cheeseboard, mean latency to reach the rewarded location as well as total path length were collected using the EthoVision software. The learning curves were normalized to percentage of 1st day latencies and distances to specifically ascertain if there were differences in the shape of the learning curves.

For the probe trial, mean distance from the reward location as well as percent time in the quadrant of the cheeseboard containing the previously rewarded location were collected.

For all exploratory tasks (Spatial, Temporal, and Sensory/Perceptual tasks), ratio values were computed after the following formula: Exploration of the object of interest (or all objects in the 5 min session of interest) minus the exploration of the other objects or last 5 min of the habituation session. This was divided by the sum of all exploration across both sessions or of both objects. As a formula this is depicted as: (A-B)/(A+B).

Exploration was defined as the mouse sniffing the object, touching the object with the paw, rearing toward the object, or whisking at the object. Touching the object with the trunk or tail or running into an object without stopping to sniff at it was not coded as exploration. Exploration was collected to the nearest .5 second.

For the reversal learning, the number of perseverative errors (continuing old rule) during the first 20 (1-20) trials were computed. The number of regressive errors (returning to old rule) were calculated during trials 21-40. A frequentist change point algorithm developed by Gallistel, Fairhurst, and Balsam (2004) and translated in the R programming language by Diep et al. (2012) was used to compute the point at which each mouse showed evidence for having learned to apply the new rule (analysis code available for download at http://github.com/mrhunsaker/Change_Point). This code takes the derivative of the learning curve at every point and determines when the slope has significantly changed. The threshold for significant change was conservatively set at p<.001 (p<.05/50) for the current task.

Data were all plotted in DataGraph (4.01 beta, Visual Data Tools, Inc. Chapel Hill, NC.). Ratio data and computed factors are plotted as bar graphs with all data points displayed. Repeated data/learning curves are presented as a line graph at the mean of each block with all data points displayed.

#### Tests for equal variance and heteroscedasticity

Prior to statistical analyses, the data were tested for normalcy (Shapiro-Wilk test) and homoscedacity (Browne-Forsythe test) to determine if the data met the assumptions for parametric analyses of variance (ANOVA). Repeated measures were evaluated for sphericity using Mauchly’s test of sphericity and necessary adjustments were made using the Huhn-Feldt correction using R 3.2.4 (Team, 2014).

#### Parametric Statistical Analysis

Once deemed appropriate, further statistical analyses were performed using parametric analyses of variance (ANOVA). For exploratory task ratios and computed factors were compared using a one-way ANOVA with groups (2N control, Ts65Dn). For acquisition tasks wherein learning was quantified across trials as well as locomotor data, statistical analyses were performed using a mixed model ANOVA with group (2N control, Ts65Dn) as a between groups factor and block of trials as a repeated within factor. An analysis was carried out comparing locomotor behaviors measured by total distance traveled on each trial in cm. In no cases were there group differences for locomotor activity (all p>.31).

All results were considered significant at an α<.05 and Power (1-β) >.80: Analyses were performed to determine observed power and effect size for all reported effects. Effect size for all analyses will be reported using the η^2^ statisic. Statistical analyses were performed in R 3.2.4 language and environment and observed statistical power was calculated using both R and the statistical program G*Power 3 (Faul, Erdfelder, Buchner, & Lang, 2009; Faul, Erdfelder, Lang, & Buchner, 2007). All reported p values were adjusted for False Discovery Rate (Benjamini, Drai, Elmer, Kafkafi, & Golani, 2001; Hunsaker, 2013) using a custom script written in R 3.2.4 (Team, 2014).

## Results

### Spatial Attribute

#### Cheeseboard

To evaluate spatial navigation and general spatial memory, mice were tested on a dry land version of the Morris water maze (cheeseboard). The Ts65Dn mice showed deficits relative to 2N control mice for raw latency to find reward (Figure 1a; groups (F(1,76)=185.645, p<.0001, η^2^=.21), no interaction among group and trial block (F(1,76)=0.333, p=.566 η^2^=.03)). These deficits are present as well when the data are adjusted for total latency on trial 1 (groups(F(1,76)=48.44, p<.0001 η^2^=.27); Figure 1b) Ts65Dn mice have impaired learning in the Ts65Dn mice in the adjusted data (F(1,76)=14.74, p=.00025 η^2^=.19). The same pattern of effects was observed for the data when evaluated for raw distance covered to find reward (Figure 1c; groups (F(1,76)=88.406, p<.0001 η^2^=.23) no interaction among group and block (F(1,76)=0.258, p=.613 η^2^=.02). Similarly to the latency data, an interaction emerges with Ts65Dn mice showing a shallower learning curve when the data are adjusted for total distance on trial 1 (groups (F(1,76)=25.194, p<.0001 η^2^=.19), interaction (F(1,76)=3.887, p=.0523 η^2^=.11); Figure 1d).

**Figure 1.**
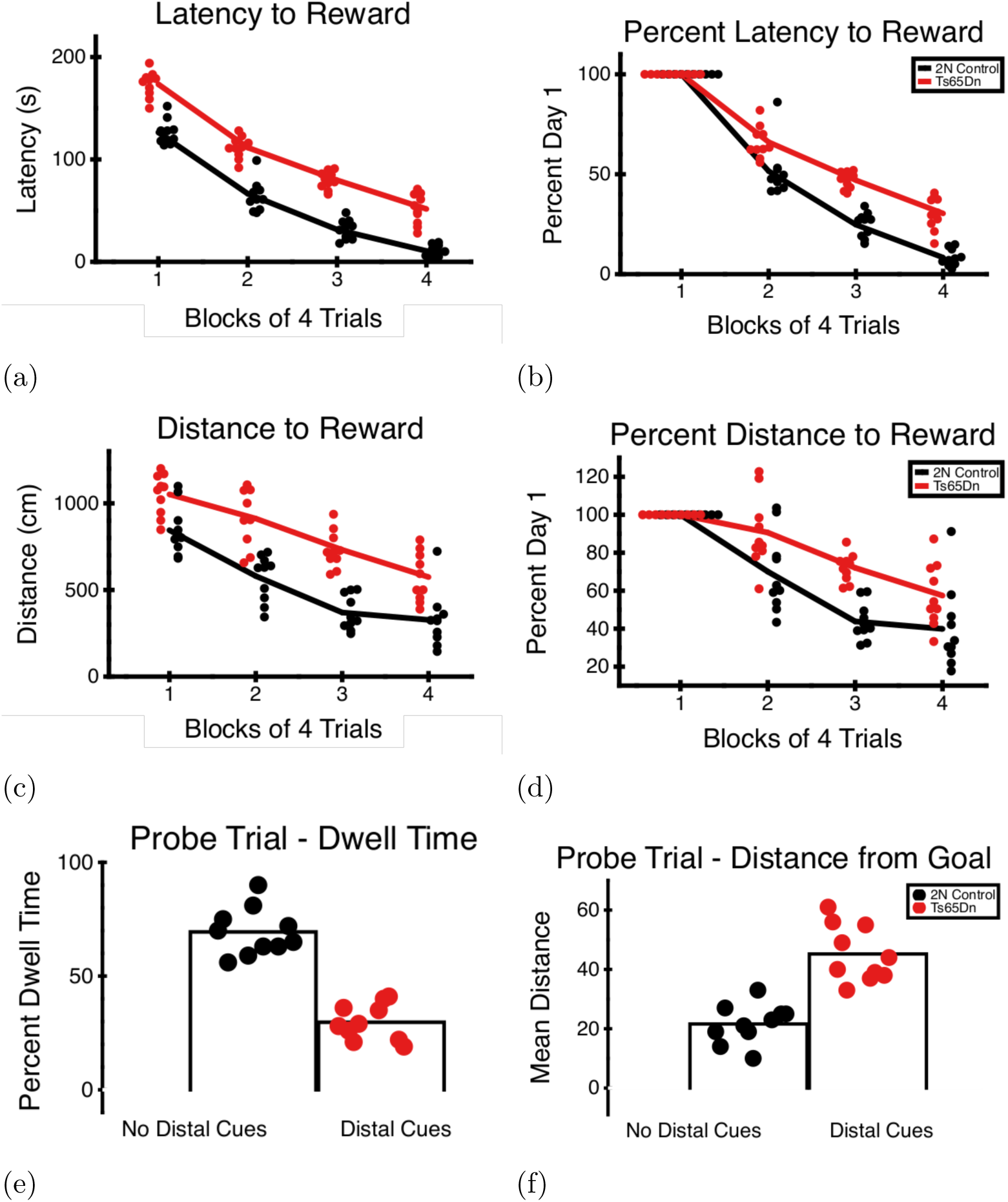
Dry land water maze performance on a cheeseboard for Ts65Dn and 2N wildtype control mice. Ts65Dn mice showed impaired spatial navigation abilities during the 4 days of acquisition, even when adjusted for initial performance. Ts65Dn mice also show spatial memory deficits during the probe trial relative to 2N wildtype control mice, reflected in reduced time in the quadrant containing the reward location and greater average distance from the previously rewarded location compared to 2N control mice. a. Raw latency (s) to reach goal location each day b. Percentage of Day 1 latency to reach goal location c. Raw distance (cm) to reach goal location d. Percentage of Day 1 distance to reach goal location e. Percentage of time during probe in same quadrant as goal location f. Average distance from goal location during probe trial

During the probe trial (Figure 1), Ts65Dn mice spent significantly less time in the quadrant where the reward was previously located (Figure 1e, F(1,18)=91.25, p<.0001 η^2^=.28). Ts65Dn mice also on average were a further distance away from the previously rewarded spatial location (F(1,18)=41.7, p<.0001 η^2^=.22; Figure 1f).

#### Metric/Coordinate processing

To evaluate coordinate / metric spatial processing, mice were tested for detection of a metric change (Figure 2a), Ts65Dn mice showed significant impairments relative to 2N control mice. There was a main effect for groups for the clear box (F(1,18)=39.38, p<.0001 η^2^=.37) as well as the red box (F(1,18)=29.94, p<.0001 η^2^=.33). Deficits in both the clear and red box suggest that metric/coordinate processing is specifically impaired in Ts65Dn mice, supporting earlier reports of dentate gyrus dysfunction in Ts65Dn mice.

**Figure 2.**
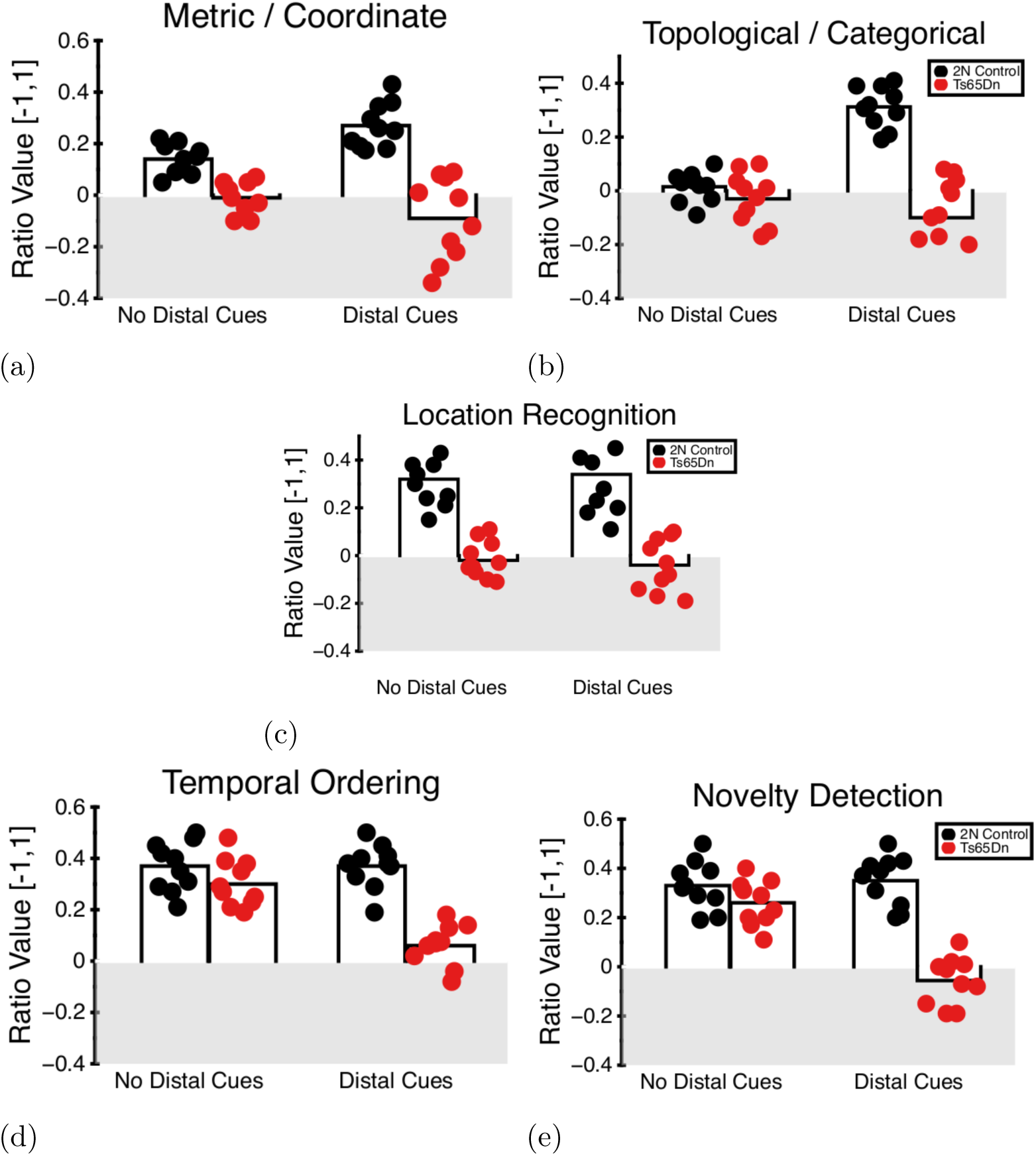
Spatial and Temporal Attribute task battery. The data suggest Ts65Dn mice show deficits relative to 2N wildtype control mice for location recognition and metric/coordinate processing, but no deficits for topological/categorical processing. The Ts65Dn mice do not show deficits for temporal ordering for visual objects compared to 2N wildtype control mice. a. Performance on a Metric / Coordinate Processing test b. Performance on a Topological / Categorical Processing test c. Performance on a Location Recognition test d.Performance on a Temporal Ordering for Visual Objects test e. Performance on a Novelty Detection for Visual Objects test

#### Topological/Categorical processing

To evaluate categorical / topological spatial processing, mice were tested for detection of a topological change (Figure 2b), Ts65Dn mice showed significant impairments relative to 2N control mice. There was a main effect for groups for the clear box (F(1,18)=78.52, p<.0001 η^2^=.24) but not for the red box (F(1,18)=1.489, p=.238 η^2^=.04). Deficits in only the clear box suggests that topological processing is only impaired when extra-maze cues are present, suggesting a general spatial memory deficit rather than one specific to topological/categorical processing.

#### Location Recognition

To test general spatial memory, mice were tested for detection of a change in the spatial location of a visual object (Figure 2c), Ts65Dn mice showed significant impairments relative to 2N control mice. There was a main effect for groups for the clear box (F(1,18)=36.39, p<.0001 η^2^=.28) as well as in the red box (F(1,18)=62.0, p<.0001 η^2^=.18), suggesting spatial novelty detection deficits in Ts65Dn mice.

### Temporal Attribute

#### Temporal Ordering of Visual Objects

To test temporal processing / temporal ordering in Ts65Dn mice, mice were tested for a simple temporal ordering task (Figure 2d). Ts65Dn mice did not show significant impairments relative to 2N control mice. There was a main effect for groups for the clear box (F(1,18)=68.24, p<.0001 η^2^=.26) but not for the red box (F(1,18)=2.267, p=.149 η^2^=.01). These data suggest that the presence of spatial cues, but not temporal ordering resulted in deficits in the clear box. For the novelty detection task run as a control for temporal ordering (Figure 2e), Ts65Dn mice did not show significant impairments relative to 2N control mice. There was a main effect for groups for the clear box (F(1,18)=82.78, p<.0001 η^2^=.21) but not for the red box (F(1,18)=2.909, p=.105 η^2^=.05). These data suggest that the presence of spatial cues, but not temporal ordering or novelty detection resulted in deficits in the clear box.

### Sensory/Perceptual Attribute

#### Feature Ambiguity

To test the ability of Ts65Dn mice to discriminate similar objects that differ only by the configuration of object features, a configural feature ambiguity test was given (Figure 3a). Ts65Dn mice did not show significant impairments relative to 2N control mice. There was a main effect for groups for the clear box (F(1,18)=34.13, p<.0001 η^2^=.,35) but not for the red box (F(1,18)=.021, p=.984 η^2^=.01). These data suggest that the presence of spatial cues, but not configural feature ambiguity resulted in deficits in the clear box. Ts65Dn mice were not impaired in a configural ambiguity control task (Figure 3b). There was a main effect for groups for the clear box (F(1,18)=12.27, p=.0025 η^2^=.15) but not for the red box (F(1,18)=.012, p=.916 η^2^=.01). These data suggest that the presence of spatial cues, but not configural feature novelty detection ordering resulted in deficits in the clear box.

**Figure 3.**
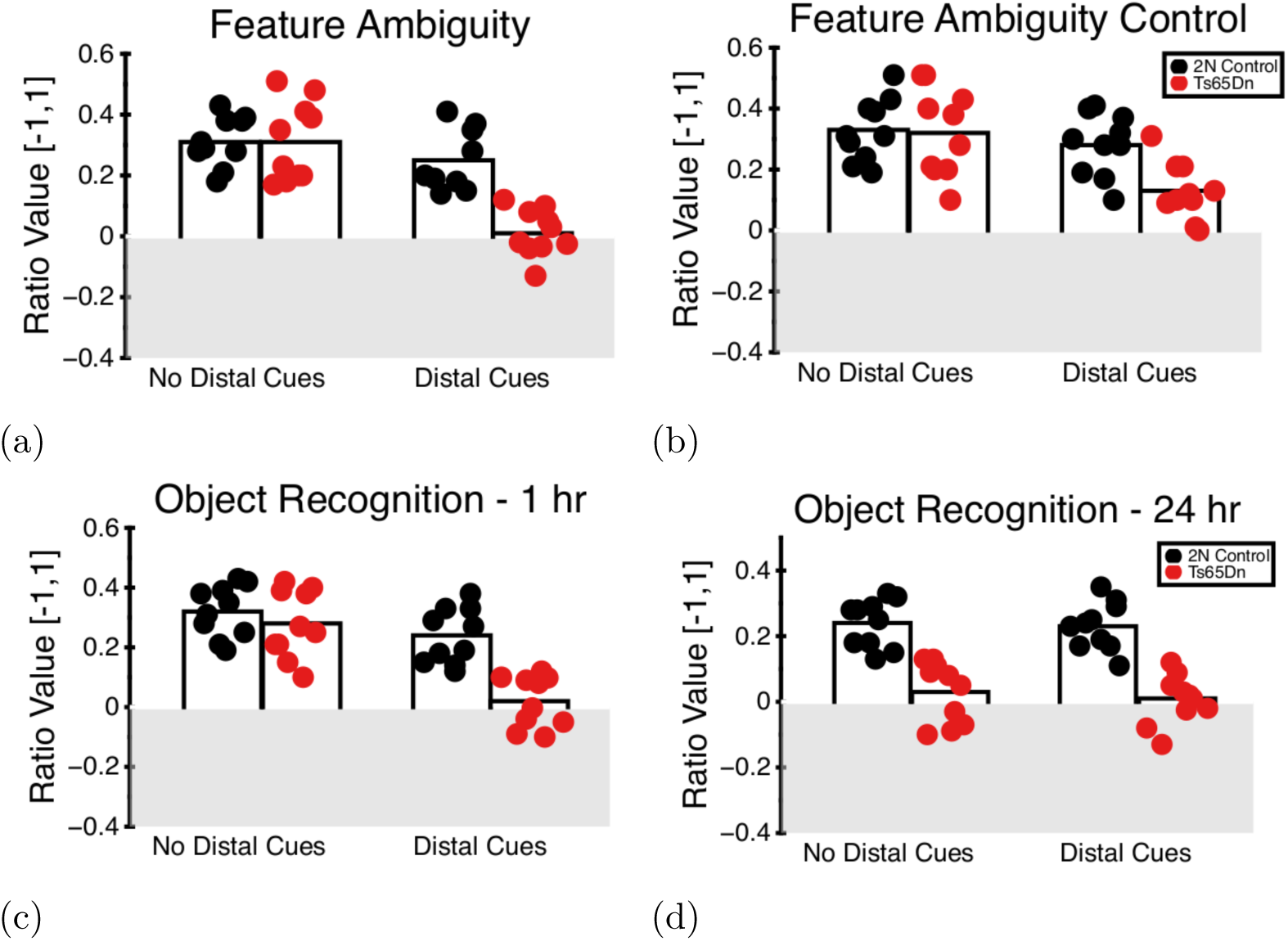
Sensory/Perceptual Attribute task battery. Overall, Ts65Dn mice do not show impaired sensory/perceptual function relative to 2N wildtype mice. Ts65Dn mice also do not show deficits for object recognition at a 1 hour delay, but do show deficits for object recognition at 24 hour delays. a. Detection of Visual Object Feature Ambiguity b. Detection of Visual Object Feature Novelty c. Performance on an Object Recognition at 1 Hour Delay test d. Performance on an Object Recognition at 24 Hour Delay test

#### Object Recognition after 1 and 24 delays

Object recognition memory was tested in Ts65Dn mice using object recognition memory at 1 and 24 hours (Figure 3c), Ts65Dn mice did not show significant impairments relative to 2N control mice. There was a main effect for groups for the clear box (F(1,18)=29.51, p<.0001 η^2^=.19) but not for the red box (F(1,18)=.908, p=.353 η^2^=.03). These data suggest that the presence of spatial cues, but not object recognition resulted in deficits in the clear box. For object recognition memory at 24 hours (Figure 3d), there was a main effect for groups for the clear box (F(1,18)=46.23, p<.0001 η^2^=.22) as well as for the red box (F(1,18)=31.36, p<.0001 η^2^=.20). These data suggest that at 24 hours, the Ts65Dn mice were unable to retrieve the memory for the object, whereas they were able to do so at 1 hour.

### Executive Function

#### Spontaneous Alternation

Spontaneous alternation was used to test working memory in the Ts65Dn mice (Figure 4a). Ts65Dn mice showed fewer alternations than 2N control mice (F(1,18)=23.85, p=.0001 η^2^=.29).

**Figure 4.**
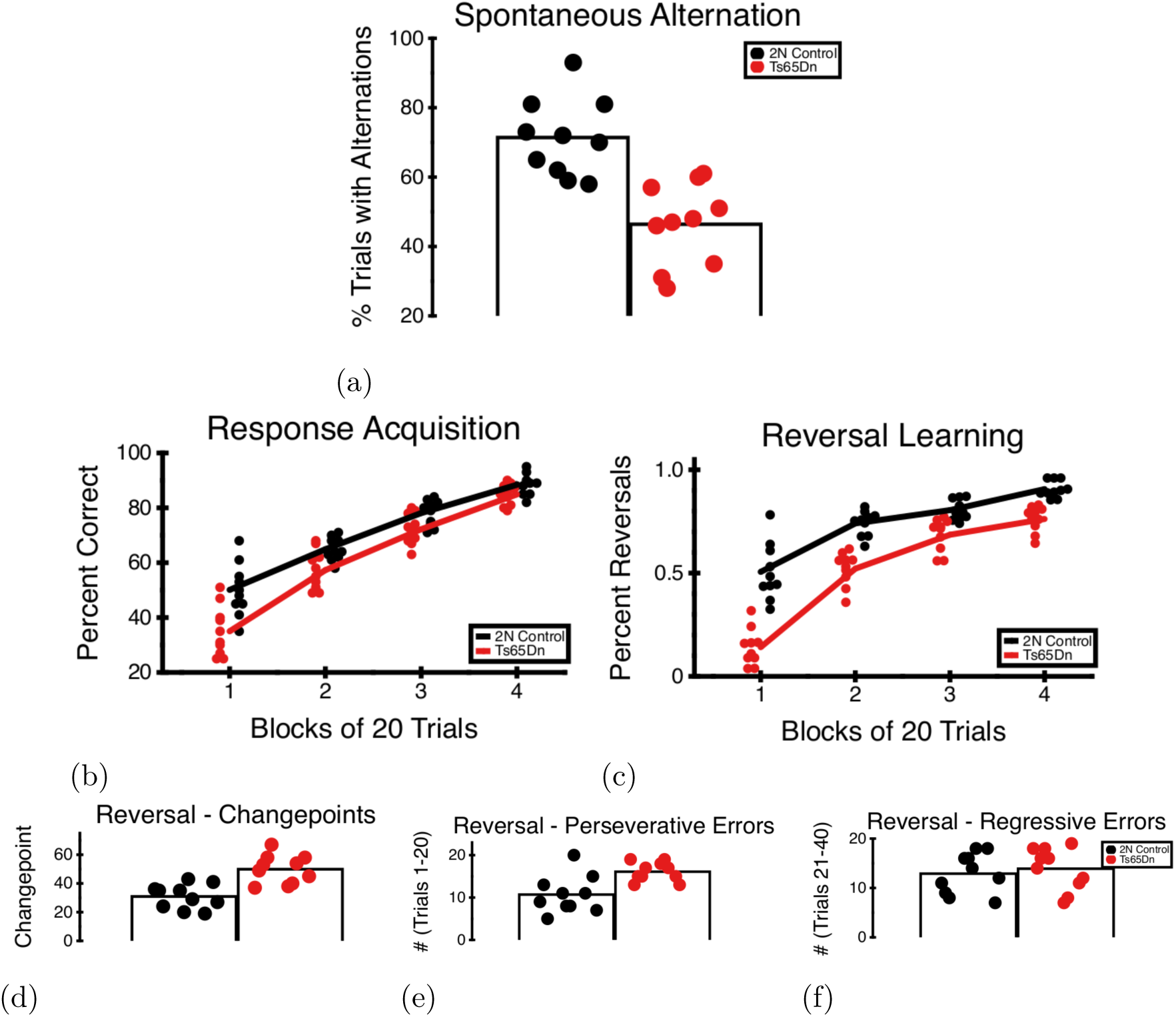
Executive Function / Rule Based Memory Task Battery. Ts65Dn mice show fewer alternations on a spontaneous alternation task relative to 2N control mice. Ts65Dn mice show mild deficits for acquisition and reversal of a rule based response on a plus maze. During reversal training, Ts65Dn mice learn to apply the new rule on later trials than control mice, reflected by an increased number of perseverative, but not regressive, errors. a. Performance on a Spontaneous Alternation test b. Acquisition of a Rule Response on a plus maze c. Acquisition of a Rule Reversal on a plus maze d. Changepoint analysis of Rule Reversal acquisition e. Perseverative Errors during trials 1-20 f. Regressive Errors during trials 21-40

#### Rule Learning on a Plus Maze

To evaluate inhibitory control and the ability to learn a turn response (Figure 4b), Ts65Dn mice took significantly longer to learn the rule than 2N control mice. There was a main effect for groups (F(1,76)=4.24, p=.013 η^2^=.14), a main effect for block of trials (F(1,76)=502.86, p<.0001 η^2^=.39). There was also an interaction among group and block (F(1,76)=7.82, p=.0065 η^2^=.23). This interaction was the result of the Ts65Dn mice taking longer to learn the rule. For the final block of 20 trials, there were no differences in performance for Ts65Dn and 2N control mice.

#### Rule Reversal Learning on a Plus Maze

To evaluate rule reversal learning (behavioral flexibility) in Ts65Dn mice, the reversal of a turn response was evaluated (Figure 4c). Ts65Dn mice took a significantly greater number of trials to learn the rule than 2N control mice. There was a main effect for groups (F(1,76)=4.952, p=.029 η^2^=.17), a main effect for block of trials (F(1,76)=24.62, p<.0001 η^2^=.17). There was also a nonsignificant interaction among group and block (F(1,76)=3.21, p=.077 η^2^=.09). Looking at Figure 4c, the nonsignificant interaction was the result of the Ts65Dn mice taking longer to learn to reverse the rule. In fact, the Ts65Dn mice were only impaired relative to the 2N control mice for the first block of 20 trials. For the remaining blocks of trials there were no differences in performance for Ts65Dn and 2N control mice. There was a main effect for groups for the trial at which the mice changed preference from old rule to new rule (changepoint; F(1,18)=21.43, p=.0002 η^2^=.13); Figure 4d). For the first 20 trials of reversal learning, Ts65Dn mice showed a greater number of perseverative errors (F(1,18)=11.98, p=.0028 η^2^=; Figure 4e). For trials 21-40, there was no difference between Ts65Dn mice and 2N control mice for regressive errors (F(1,18)=.287, p=.599 η^2^=.02; Figure 4f).

### Motor Function

#### Capellini Eating Task

For the capellini task of manual dexterity (Figure 5), Ts65Dn mice showed significant impairments relative to 2N control mice. There was a main effect for latency, with Ts65Dn mice taking longer to eat the pasta on average (F(1,18)=14.74, p=.0012 η^2^=.17; Figure 5a). Ts65Dn mice also made a greater number of pasta handling errors (F(1,18)=92.68, p<.0001 η^2^=.40; Figure 5b). There was also a main effect for groups for the number of times the paws came together (F(1,18)=42.34, p<.0001; Figure 5c), for the number of times the mouse lost contact with the pasta (F(1,18)=20.35, p=.0003 η^2^=.22; Figure 5d) and the number of times the mouse pulled the pasta with their mouth rather than using the hands to move it (F(1,18)=21.46, p=.0002 η^2^=.17; Figure 5e).

**Figure 5.**
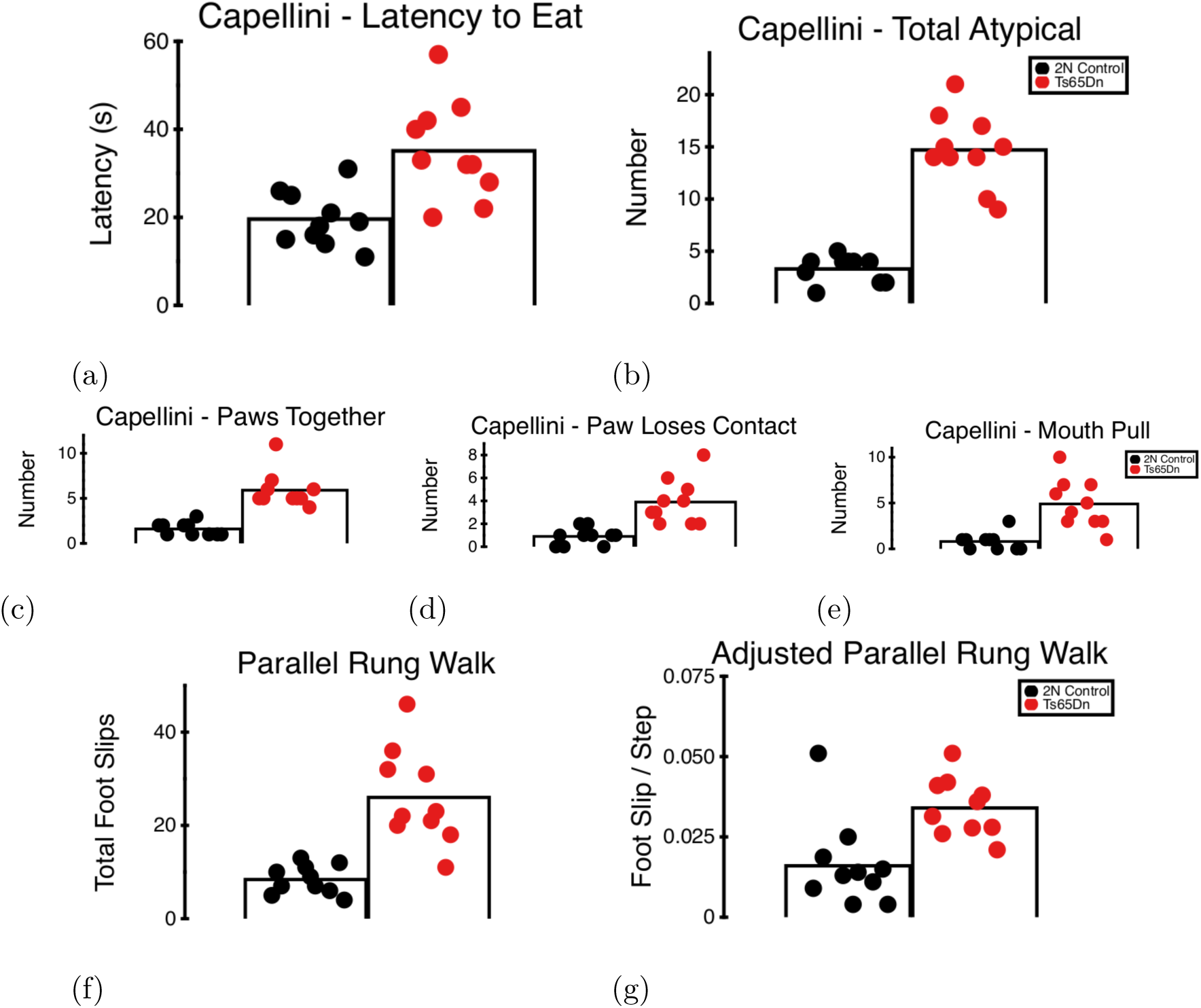
Motor Function Task Battery. Ts65Dn mice showed reduced motor dexterity during a Capellini Handling task reflected as an increase in the number of abnormal behaviors and increased latency to consume the capellini as well a greater number of foot slips during a Parallel Rung Walking task, even when adjusted for total number of steps. a. Latency (s) to consume capellini b. Total number of abnormal behaviors c. Number of times paws came together and touched d. Number of times paw lost contact e. Total number of times mouth was used to move capellini f. Total number of foot slips on a Parallel Rung Walking test g.Total number of foot slips when adjusted for total number of steps

#### Parallel Rung Walking Task

During a parallel rung walking task (Figure 5f), Ts65Dn mice showed significant impairments relative to 2N control mice. There was a main effect for the number of foot slips in a 1 minute session (F(1,18)=27,32, p<.0001 η^2^=.19). When adjusted for number of steps, Ts65Dn mice still showed a greater number of foot slip errors (F(1,18)=11.70, p=.0031 η^2^=.16; Figure 5g).

### Adaptive Function / Quality of Life

#### Nesting Behavior

Ts65Dn mice showed significant impairments relative to 2N control mice for measures of nesting (Figure 6). Ts65Dn mice took longer to make contact with the nesting material (F(1,18)=152.9, p<.0001 η^2^=.24; Figure 6a), for the time it took for them to dig in the media (measured from time of first contact) (F(1,18)=318.6, p<.0001 η^2^=.16; Figure 6b), and the time it took from starting to dig to finish the nest (F(1,18)=94.3, p<.0001 η^2^=.21; Figure 6c).

**Figure 6.**
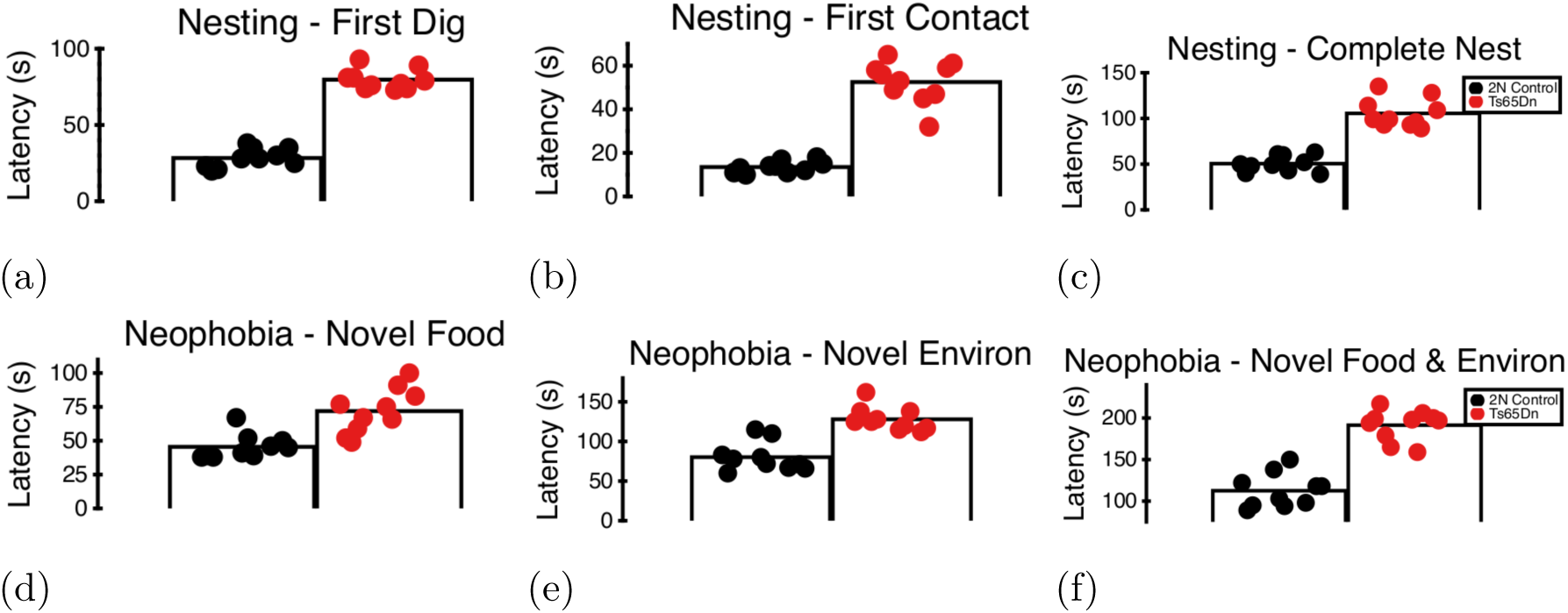
Adaptive Function / Quality of Life Task Battery. Ts65Dn mice take longer to make a nest out of preferred nesting material and show increased neophobia for both food and environments. a.Latency (s) to initially contact nesting material. b. Latency (s) to begin digging in nesting material c. Total latency (s) to finish nest d. Latency (s) to begin consuming novel food in familiar environment e. Latency (s) to consume familiar food in novel environment. f. Latency (s) to consume novel food in novel environment

#### Neophobia

Ts65Dn mice showed significant impairments relative to 2N control mice for neophobia (Figure 6). Ts65Dn mice took longer to eat a novel food in a familiar environment (F(1,18)=19.59, p=.0003 η^2^=.11; Figure 6d), took longer to eat a familiar food in a novel environment (F(1,18)=40.87, p<.0001 η^2^=.16; Figure 6e), and took longer to eat a novel food in a novel environment (F(1,18)=83.74, p<.0001 η^2^=.17; Figure 6f).

## Discussion

Briefly, Ts65Dn mice displayed specific deficits for spatial processing, long-term memory, motor function, executive function, and adaptive function (Table 1). These deficits phenocopy the results from the ACTB used in testing children with Down syndrome, including the report that providing distracting contextual cues may impair memory function in Down syndrome (Edgin et al., 2010; Edgin et al., 2012; Edgin et al., 2014).

Overall, these data clearly demonstrate that the Ts65Dn mouse do in fact show a similar pattern of behavioral deficits on the mouse variant of the Arizona Cognitive Task Battery (mCTB) as individuals with Down syndrome show on the human ACTB. The task similarities between the mouse and human ACTB are outlined in Table 1. In cases where Down syndrome participants show deficits on the ACTB (Edgin et al., 2010), the mice in the present study phenocopy those effects (also *cf.*, Edgin et al. (2012)). Similarly. the Ts65Dn mice showed the same pattern of strengths (*i.e.*, lack of performance deficits) as individuals with Down syndrome show on the ACTB.

The pattern of Ts65Dn performance on spatial and temporal processing tasks support the hypothesis that Ts65Dn mice show clear deficits for spatial processing tasks dependent upon the dentate gyrus with sparing of spatial and temporal processing dependent upon the CA1 subregion (Goodrich-Hunsaker, Hunsaker, & Kesner, 2008; Kesner, Lee, & Gilbert, 2004; Kesner & Rolls, 2015; Rolls & Kesner, 2006; Smith et al., 2014). Similarly, it appears that spatial processing dependent on neocortical processing is spared (*cf.,* Goodrich-Hunsaker, Hunsaker, and Kesner (2005)). Similar cognitive deficits have been reported in Down syndrome (Edgin et al., 2012).

These findings were confirmed by verifying that any spatial or temporal processing deficits observed in the presence of distal cues was confirmed in a task that removed these cues (Dees & Kesner, 2013). The data show that metric/coordinate processing and location recognition deficits are similar in the presence or absence of distal cues, suggesting that these hippocampus (more specifically the dentate gyrus) dependent spatial processes are disrupted. The topological/categorical deficits observed in the clear box are absent when tested in the absence of extramaze cues in a red box. These data suggest that CA1/parietal cortex related spatial memory processes are intact when tested without extra-maze cues available (*cf.*, Kesner et al. (2004), Kesner and Rolls (2015)).

Similarly, the temporal ordering deficits present in the clear box were absent in the red box, and the novelty detection control task showed the same pattern, suggesting temporal processing is intact in the Ts65Dn mice, but object identification may be impaired if extra-maze distal cues are present. This hypothesis was confirmed in the sensory/perceptual tests wherein the Ts65Dn mice were able to correctly process feature ambiguity and feature novelty in the red, but not clear boxes. And finally, object recognition was impaired even at only 1 hour delays for Ts65Dn mice when extramaze cues were available. In the red box, the Ts65Dn mice were able to identify previously encountered objects until a 24 hour delay was imposed.

For response learning or executive function, Ts65Dn mice were impaired for spontaneous alternation (they alternated on fewer trials than wildtype mice), as well as response learning and reversal learning of a previously learned rule. However, it appeared that the Ts65Dn mice just learned the tasks more slowly since the early trials show deficit, but later blocks of trials do not. For reversal learning, it is clear the Ts65Dn mice take a greater number of trials to learn the reversal based on the changepoint calculated for the learning curves (Ts65Dn mean=50 compared to mean=30 for 2N wildtype mice) as well as the greater number of perseverative errors during trials 1-20 of the reversal learning task. Interestingly, once the Ts65Dn mice showed learning of the reversal, they did not make any more regressive errors than the 2N control mice.

These data support earlier theories that suggested there were specific deficits to spatial memory in Down syndrome (Carlesimo, Marotta, & Vicari, 1997; Carretti & Lanfranchi, 2010; Lanfranchi et al., 2009; Silvia Lanfranchi et al., 2004; Vicari et al., 2005; Visu-Petra, Benga, Miclea, et al., 2007). What these data clarify are the neural substrates and specific domains of medial temporal lobe function are impaired in Down syndrome. There are specific deficits on tasks that test dentate gyrus function, but sparing of function on tasks that test parietal and perirhinal cortices as well as CA1 function. Similarly, there are specific deficits in the Ts65Dn mouse that are attributable to cerebellar function and executive functional deficits attributable to the rostral cortices (analogue of the human prefrontal cortex). For thorough descriptions of neuroanatomic correlates of the behavioral tasks included in the mCTB the reader is referred to the descriptions of the original tasks (*cf.*, Bartko et al. (2007), Bussey et al. (2002), Kesner et al. (2004), Kesner and Rolls (2015), Ragozzino et al. (1999), Ragozzino et al. (2002), Rolls and Kesner (2006)

For the motor tasks, the Ts65Dn mice showed clear deficits for handling the capellini and greater difficulties walking on parallel rungs. For adaptive function, the Ts65Dn nice took longer to build nests and consume novel foods in novel locations, suggesting reduced adaptive function or quality of life relative to 2N control mice.

An important consideration in adopting a behavioral screen like this mCTB is the relative throughput for the tasks. All of the tasks used to test medial temporal lobe function take 30 minutes per session of testing, and can be repeated numerous times on any given mouse after 24 hours have passed since the first test. The motor and adaptive function tests are similarly high throughput, as is the spontaneous alternation task. The only tasks that require a significant time investment are the dry land watermaze (Lopez et al., 2010) on the cheeseboard and the rule acquisition and rule reversal learning tasks (Bissonette et al., 2008; Ragozzino et al., 1999; Ragozzino et al., 2002). The dry land watermaze task on the cheeseboard follows a standard water maze protocol that lasts 5 days, and the response learning and reversal learning tasks together take an additional week.

A second consideration is adopting the mCTB is the advantage of the anatomical specificity of known neural substrates underlying each behavioral task (Bartko et al., 2007; Bussey et al., 2002, 2006; Farr et al., 2006; Goodrich-Hunsaker et al., 2005, 2008; Hunsaker, 2012a; Kesner et al., 2004; Kesner & Rolls, 2015) and previous comparison of rodent performance on many of the behavioral tasks to human cognitive function (Baumann, Chan, & Mattingley, 2012; Baumann & Mattingley, 2013; Goodrich-Hunsaker & Hopkins, 2010; Goodrich-Hunsaker et al., 2005; Kesner & Goodrich-Hunsaker, 2010). As such, these tasks can be used to dissociate function of brain areas within the mouse models being tested. The final consideration is the lack of negative reinforcement or aversive stimulus. This means mouse models displaying depression, anxiety, or anhedonia are theoretically testable using the mCTB (*cf.*, Hunsaker (2012a, 2012b)).

An interesting complication emerged in the data that the mCTB was solved by nature of how it was designed. On a number of nonspatial tasks. there was a confound of distal cues interfering with the processing of proximal objects that were of interest in the task. For example, in the temporal ordering and novelty detection for novel objects tasks, the Ts65Dn mice looked like they had deficits, but only in the clear box that allowed access to distal cues (Dees & Kesner, 2013; Smith et al., 2014). The feature ambiguity task and the control condition showed the same pattern. The addition of a distal cue-free condition (the red box) was essential for separating the effects of proximal-distal cue interactions from the memory processes being tested by the tasks. The disparate performance across clear and red boxes (or in presence of absence of extra maze contextual cues) allowed us to assess the role of context and distracting cues in memory function in Ts65Dn mice, a conceptual replication of Edgin et al. (2014) in Down syndrome and rats as shown by Dees and Kesner (2013).

## Limitations

The primary limitation of the present study is the lack of tests for language or language like attributes in the Ts65Dn mouse model. However, such assays exist and can easily be added to the task battery without significantly increasing the amount of time required to perform the mCTB (Zampieri, Fernandez, Pearson, Stasko, & Costa, 2014). The present experiment also only assayed the Ts65Dn mouse model of Down syndrome as a proof of concept. Further studies will be necessary to evaluate whether other mouse models of Down syndrome (*e.g.,* Ts2Cje, Ts1Yah, and Dep(17)1Yey/+; Das and Reeves (2011)) show the same pattern of results as the Ts65Dn mouse model. This is not a trivial issue as there is still controversy as to which of the many genetic models best recapitulate the cognitive phenotype seen in Down syndrome populations.

## Conclusions

That deficits in the mouse and human ACTB are comparable suggests that the mCTB may be useful for guiding the development of treatment strategies by providing reliable, valid behavioral endpoints and outcome measures. These outcome measures reported in the mCTB appear to show high face, content, and predictive validity with the ACTB, at least so far as Ts65Dn performance mimics the performance of Down syndrome patient populations. As we were able to identify such a clear phenotype in Ts65Dn mice, the mouse mCTB may well turn out to be a useful tool for studying behavioral prodrome of early Alzheimer-like pathology and cognitive decline in mouse models related to Down syndrome. Similarly, the mCTB may serve as a powerful and comprehensive screening tool for preclinical tests of pharmacological interventions in Down syndrome.

